# Repeated passive visual experience modulates spontaneous and non-familiar stimuli-evoked neural activity

**DOI:** 10.1101/2023.02.21.529278

**Authors:** Suraj Niraula, William L. Hauser, Adam G. Rouse, Jaichandar Subramanian

## Abstract

Familiarity creates subjective memory of repeated innocuous experiences, reduces neural and behavioral responsiveness to those experiences, and enhances novelty detection. The neural correlates of the internal model of familiarity and the cellular mechanisms of enhanced novelty detection following multi-day repeated passive experience remain elusive. Using the mouse visual cortex as a model system, we test how the repeated passive experience of a 45° orientation-grating stimulus for multiple days alters spontaneous and non-familiar stimuli evoked neural activity in neurons tuned to familiar or non-familiar stimuli. We found that familiarity elicits stimulus competition such that stimulus selectivity reduces in neurons tuned to the familiar 45° stimulus; it increases in those tuned to the 90° stimulus but does not affect neurons tuned to the orthogonal 135° stimulus. Furthermore, neurons tuned to orientations 45° apart from the familiar stimulus dominate local functional connectivity. Interestingly, responsiveness to natural images, which consists of familiar and non-familiar orientations, increases subtly in neurons that exhibit stimulus competition. We also show the similarity between familiar grating stimulus-evoked and spontaneous activity increases, indicative of an internal model of altered experience.

## Introduction

Long-term repeated experience of an innocuous stimulus leads to behavioral habituation and a subjective memory of the stimulus. Habituation enables organisms to respond to novel or behaviorally relevant stimuli^1,2^. Repeated viewing of an orientation-grating not associated with reward or punishment over multiple days selectively reduces behavioral exploration of that orientation in mice^3,4^. However, at the level of neural activity, the effect of the multi-day repeated experience of orientation grating stimuli has yielded contradictory findings. Studies have found a familiar stimulus-selective increase in local field potential and altered oscillations^5-9^. Single-unit recordings have found an increase in peak responsiveness and a decrease in average responsiveness to familiar grating stimuli^3,10^. Calcium imaging studies have yielded mixed results with no change^11^, increased^12^, or decreased average responsiveness^13-15^ to familiar grating stimuli. Our recent study found that familiarity with a grating stimulus reduces average neural responsiveness to that stimulus, consistent with some earlier studies^4^. While the reduction in average responsiveness to the familiar stimulus could explain behavioral habituation, how repeated passive non-natural stimuli experience over multiple days alters spontaneous activity and enhance novelty detection at the cellular level is unclear.

Functionally distinct inputs, including those tuned to different orientations, are intermingled on dendrites of visual cortical neurons^16-19^. Therefore, reduced responsiveness to the familiar stimulus could trigger competition and enhance responsiveness to novel stimuli. Short-term visual adaptation to an oriented grating or Gabor stimulus shifts the tuning curve toward or away from the adapted orientation depending on various stimulus features and enhances novelty detection^20-29^. Similarly, the multi-day repeated passive experience of orientation grating stimulus has been shown to increase orientation selectivity^10^, whereas other studies found a lack of orientation tuning curve shift^13-15^. Whether multi-day passive experience enhances long-term (>12 hours) responsiveness to non-familiar stimuli and whether it involves competition between familiar and non-familiar stimuli remains unclear.

Expectations or subjective memory of the familiar stimulus may arise from a stored internal model. Spatiotemporal characteristics of spontaneous activity, which occurs without stimulus, resemble evoked activity and may serve as an internal model of an animal’s sensory environment^30-42^. However, recent studies in mice and zebrafish show that spontaneous and visually evoked activity are dissimilar, further diverging over development ^43-45^. If spontaneous activity serves as an internal model of sensory experience, the repeated passive sensory experience should increase the similarity of spontaneous and familiar experience-evoked activity patterns in adulthood.

We show that in layer 2/3 of the mouse visual cortex, familiarity with a grating stimulus enables competitive plasticity between repeatedly experienced (familiar) and once experienced (non-familiar) orientations leading to higher responsiveness to some of the non-familiar stimuli (cardinal orientations and natural images) than when these stimuli were novel. Altered responsiveness to familiar and non-familiar stimuli changes stimulus selectivity differentially dependent on the stimulus preference of neurons. Interestingly, the fraction of neurons activated by natural images increased subtly, and the neurons that became more responsive to natural images exhibited higher stimulus competition than those that became less responsive. Furthermore, spontaneous and familiar stimulus-evoked activity patterns became more similar and shared the same neural space. Based on these observations, we speculate that superficial visual cortical neurons store an internal model of altered experience by increasing the similarity between spontaneous and familiarity-evoked activity and detect deviation from familiarity through changes in stimulus selectivity elicited by stimulus competition.

## Results

### Repeated passive exposure to specific orientation reduces neural responsiveness selectively to the familiar stimulus

We imaged spontaneous and visually evoked calcium transients in the awake head and body-restrained GCaMP6s transgenic mice habituated to the imaging apparatus (Fig. 1A, B). Visual stimuli consisted of eight trials of sinusoidal phase-reversing orientation gratings and a set of ten natural images (Fig. 1B). The phase-reversing orientations do not drift, thus 0°, 45°, 90°, and 135° are the same as 180°, 225°, 270°, and 315° (hereafter referred to as 0°,45°,90°, and 135°, respectively). For the next eight days, head-fixed mice passively experienced two sessions of 60 seconds of the gray screen followed by five blocks of 100 seconds of 45° phase reversing grating stimulus with 30 seconds of the gray screen between blocks. The two sessions were separated by ∼1-2 hours. We refer to the passive repeated exposure to the non-natural grating stimulus for eight days as an “altered experience.” The next day after the altered experience, the same neurons were imaged, similar to the first imaging session (Fig. 1B-F). 1375 neurons from ten mice (7 males and 3 females) were identified from both sessions combined and ranked based on the quality of soma in the anatomical and correlation maps generated by Suite2p^46^. 561 neurons with clearly visible somata in both sessions were included for further analyses.

**Figure 1.**
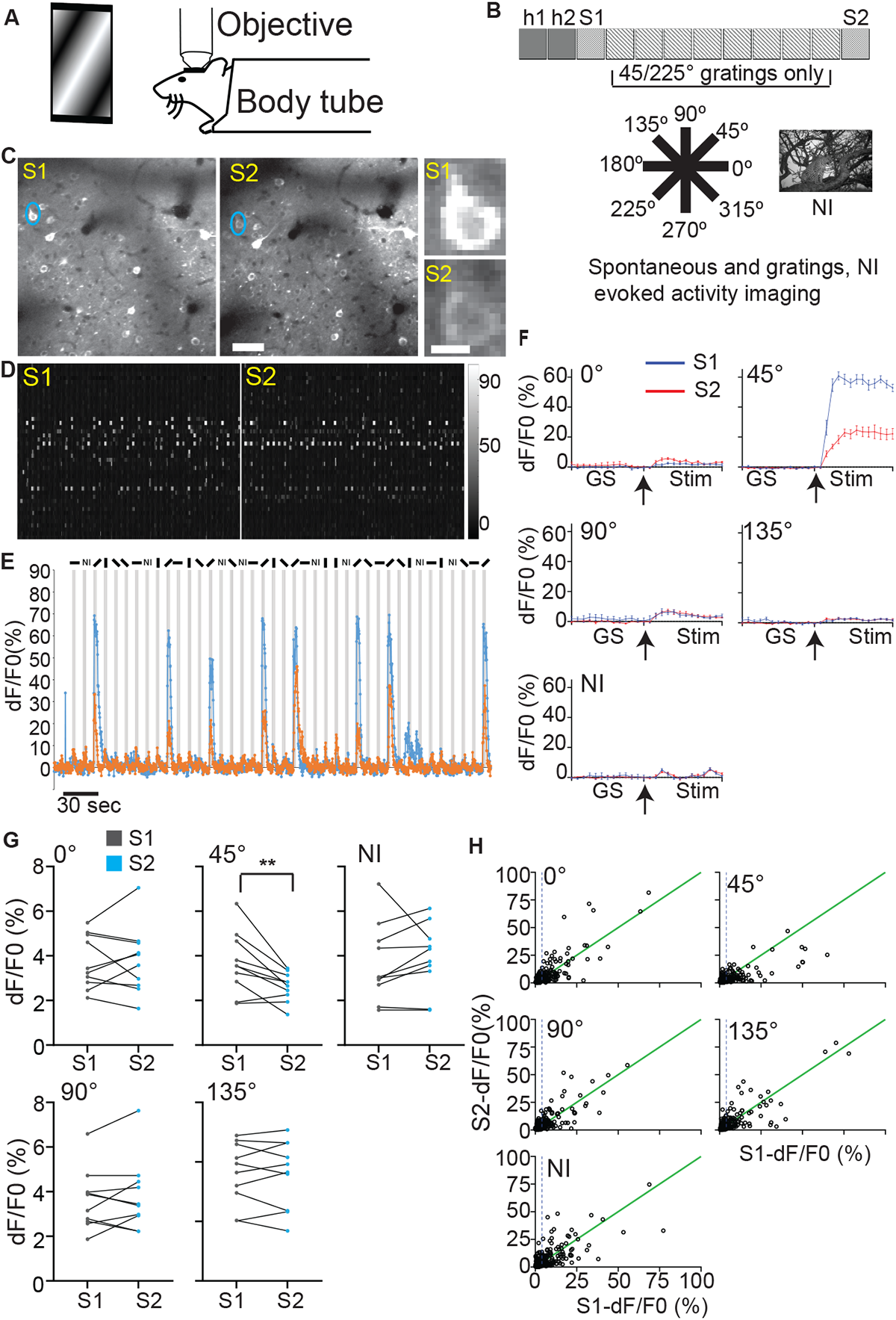
Repeated orientation-grating stimulus exposure reduces responsiveness selectively to the familiar stimulus. **A.** representation of a head and body restrained mouse placed under microscope objective viewing visual stimuli. **B.** Experimental timeline (top). Each square indicates a day, and h1 and h2 are habituation days when mice were exposed to a gray screen. Spontaneous and visually evoked calcium transients were imaged in sessions S1 and S2. Visual stimuli consisted of full-field phase reversing grating stimuli of different orientations and natural images (NI; bottom). On the intervening days, mice experienced only one orientation of phase reversing grating stimulus. **C**. Representative standard deviation projection images from S1 (left) and S2 (right); scale bar: 50 μm. A neuron circled in blue is zoomed on the right; scale bar: 10 μm. **D.** Raster plot of calcium transients (dF/F0 indicated by the color bar) from identified neurons in the imaging field in C on S1 (left) and S2 (right). **E.** dF/F0 of calcium transients from the neuron circled in C on S1 (blue) and S2 (orange) during the entire imaging period encompassing gray screen, different grating, and natural images (represented as bars or NI on top) stimuli. Gray bars represent the stimulus duration. **F.** Trial-averaged dF/F0 (%) of the transients in E for each stimulus. GS and stim (3 seconds each) represent the gray screen and the indicated stimulus period, respectively. Arrow represents the start of the stimulus. Data presented as mean ± SEM. **G**. Population average of the mean trial-averaged dF/F0 for the indicated stimulus on S1 and S2. ** *p*<0.01; paired *t* test; *n* **=** 10 mice. Circles represent the average for each mouse and the lines connect the values from the same mouse in the two sessions**. H.** Scatterplot representing the relation between dF/F0 on S1 and S2. The circles represent neurons (561 neurons). The dotted line represents the approximate mean dF/F0 on S1 (4%) and values less and more than the mean are considered low and highly responsive neurons, respectively. The green line represents the identity line.

To compare visually evoked calcium transients, we matched neurons across imaging sessions pre (S1) and post (S2) altered experience. We found that the trial-averaged mean dF/F0 over the 3-second stimulus period from all neurons matched between sessions selectively reduced to 45° (familiar) but not to other grating or natural image stimuli (non-familiar) (Fig. 1G). Familiar stimulus-specific reduction in population response indicates that plasticity is not an artifact associated with the animal’s behavioral state.

We found that the average dF/F0 elicited by all stimuli are significantly correlated between sessions; however, the response of above-average responsive neurons lay farther from the identity line, indicating highly responsive neurons are more dynamic, with large increases and decreases from their initial dF/F0 (Fig. 1H). In contrast to non-familiar stimuli, highly responsive neurons to the familiar 45° stimulus mostly showed a reduction in dF/F0 (Fig. 1H).

### Familiarity alters the tuning preference of low but not high-responsive neurons

Due to the intermingling of synapses with multiple orientation preferences, even in neurons selectively tuned to one of the orientations, a reduction in responsiveness to familiar orientation is likely to increase the responsiveness to non-familiar orientations. Such competition underlies short-term adaptation to orientation grating responsiveness^29^. We now tested, whether long-term (>12 hours) changes to preferred orientation also happen following multi-day grating stimulus exposure. Stimulus competition may influence tuning properties differently depending on neuronal activity levels. A small shift in responsiveness to familiar and non-familiar orientations may alter the preferred orientation for low but not highly responsive neurons. First, we classified neurons as stimulus responsive if the trial averaged mean dF/F0 is greater than two standard deviations above the mean trial averaged gray screen period preceding the stimulus. We next classified them as high and low-responsive neurons if the mean of the trial-averaged dF/F0 is greater and lower than the population’s mean dF/F0 (∼4% for all stimuli), respectively.

We next identified neurons that were classified as responsive with good tuning curve fit in both sessions (286/561 neurons; Fig. 2A). We binned these well-fit neurons by their preferred orientation into four groups centered on the four stimuli (0° – 22.5° and >157.5°-180° as 0°, >22.5°-67.5° as 45°, >67.5°-112.5° as 90°, and >112.5°-157.5 as 135° tuned neurons). We found a modest reduction in neurons whose preferred orientation was closer to the familiar orientation (∼27, 23, 26, and 24% of neurons were tuned to 0 °,45 °,90 °, and 135°, respectively, in S1 but changed to 31, 15, 28, and 26% for these orientations; Fig. 2B, top). However, when restricting this comparison to neurons that were highly responsive in both sessions, the proportion of neurons tuned to different orientations did not differ (Fig. 2B, middle). In contrast, the distribution of tuning preferences differed for neurons that were low responsive in both sessions (31, 23, 24, and 23% for 0 °,45 °,90 °, and 135°, respectively in S1 and 36, 11, 30, and 22% for these orientations in S2; Fig. 2B, bottom). These results indicate that altered experience shifts the tuning preference of neurons that were low but not highly responsive in both sessions. We next compared the fraction of neurons that retained their original tuning preference after the altered experience. When we compared the neurons responsive in both sessions with good tuning curve fit, we found that 73% of neurons maintained their tuning preference within their bin if the preferred stimulus did not become familiar, compared to 56% if it became familiar (Fig. 2C, D, top). Consistent with our observation that the distribution of tuning preferences did not change for neurons that retained their high responsiveness between sessions, we found that a comparable fraction of highly responsive neurons retained their original stimulus preference regardless of familiarity (82% of neurons preferring non-familiar and 71% of neurons preferring familiar stimulus retained their tuning preference; Fig. 2C, D, middle). In contrast, 38% of neurons that were low responsive in both sessions retained their preferred orientation when the preferred stimulus became familiar, whereas 62% retained it when it remained non-familiar (Fig. 2C, D, bottom). These results suggest that familiarity elicits competition between stimuli, so non-familiar stimulus responsiveness becomes more dominant in neurons that stayed low-responsive pre- and post-altered experiences.

**Figure 2.**
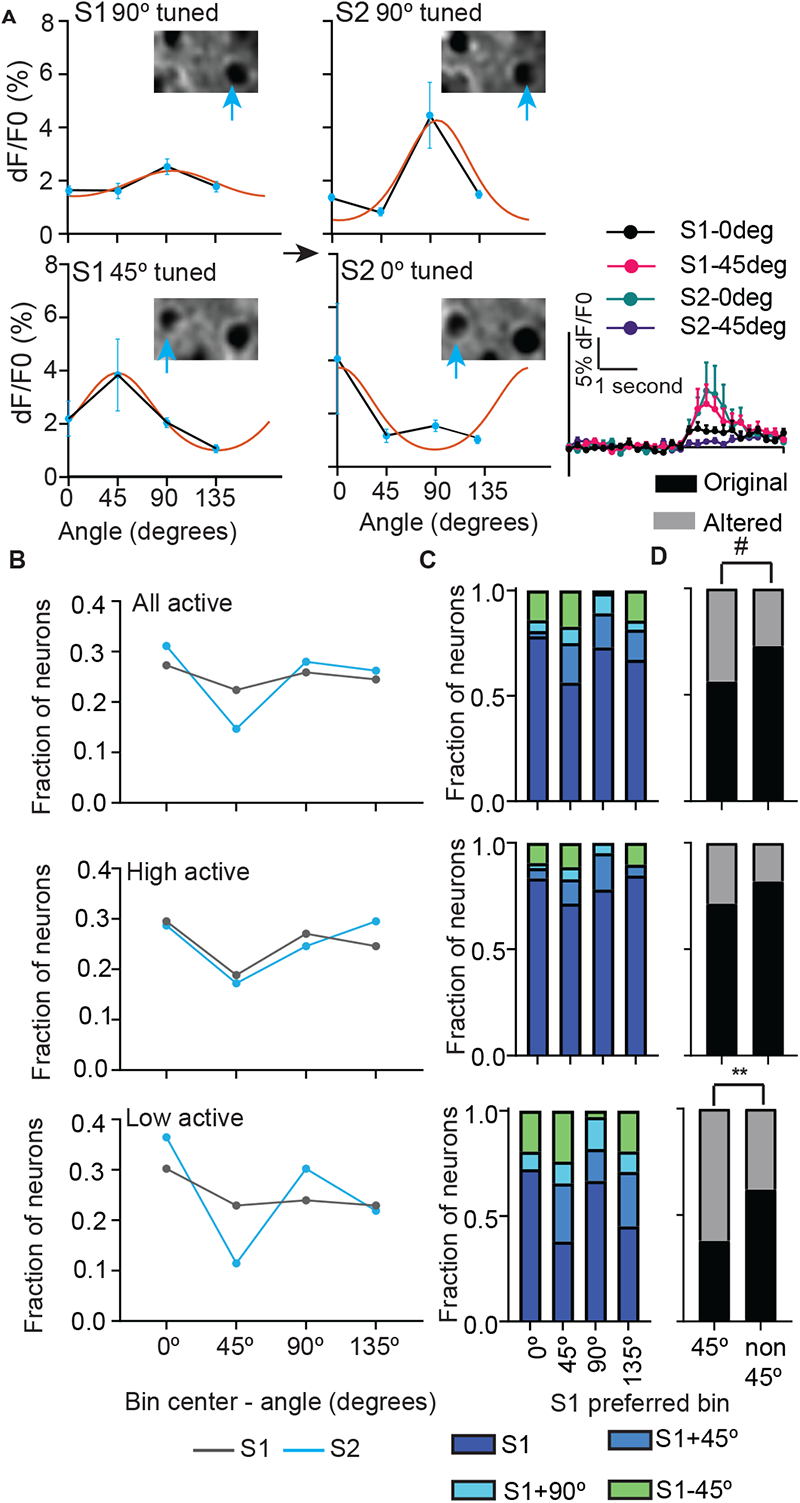
Altered experience elicits stimulus competition in low responsive neurons. **A.** Representative orientation tuning curve fit (orange line) of a low responsive neuron that retained (top) or shifted (bottom) its preferred stimulus following altered experience. The blue circles represent the mean of trial-averaged dF/F0 of 0°, 45°, 90°, and 135° grating stimuli, and error bars are SEM for the 8-trials of representative neurons (indicated by the blue arrow on the images at the top). Bottom right – Trial averaged dF/F0 trace elicited by 0° and 45° stimuli to represent the change in the preferred stimulus. **B.** fraction of all (top; 286 neurons), high (middle; 122 neurons), or low (bottom; 96 neurons) grating responsive neurons with good tuning curve fit in both imaging sessions in indicated bins separated by 45° before (S1) and after (S2) altered experience. **C.** fraction of all (top), high (middle), or low (bottom) neurons tuned to indicated S1 orientation that retained their tuning within the same bin or changed (± 45° or + 90° relative to S1). **D.** Same as C except that 0°, 90°, and 135° (non-45°) bins are pooled for all (top; n = 222 non-45° and 64 45° neurons), high responsive (middle; n = 122 non-45° and 35 45° neurons), and low (bottom; 100 non-45°, 29 45° neurons) responsive neurons. Original and not original represents the fraction of neurons that retained and changed their S1 preference, respectively. # *p* = 0.05, ** *p* < 0.01 McNemar’s test for correlated proportions.

### Familiarity differentially alters the tuning width of highly responsive neurons based on their tuning preference

To test whether stimulus competition occurs in neurons that retained highly responsiveness to grating stimuli following altered experience, we compared the tuning width of these neurons (Fig. 3A). Decreased responsiveness to the familiar stimulus or increased responsiveness to non-familiar stimulus would broaden and sharpen the tuning width of neurons tuned to familiar or the same non-familiar stimulus, respectively. We found that the median full width at half-maximum was ∼ 59°, slightly broader than the 22-29° half-width at half-maximum reported previously^47^. Altered experience differentially influenced the tuning width of neurons tuned to familiar and some non-familiar stimuli (Fig. 3B-E). Neurons tuned to 0° showed a non-significant left shift of tuning width distribution, and 135° (orthogonal to familiar stimulus) had no change (Fig. 3B, E). In contrast, 90° tuned neurons showed a significant left shift of frequency distribution and reduced average tuning width, whereas familiar stimulus-tuned neurons showed a right shift (not significant) and a subtle increase in average tuning width (Fig. 3C, D).

**Figure 3.**
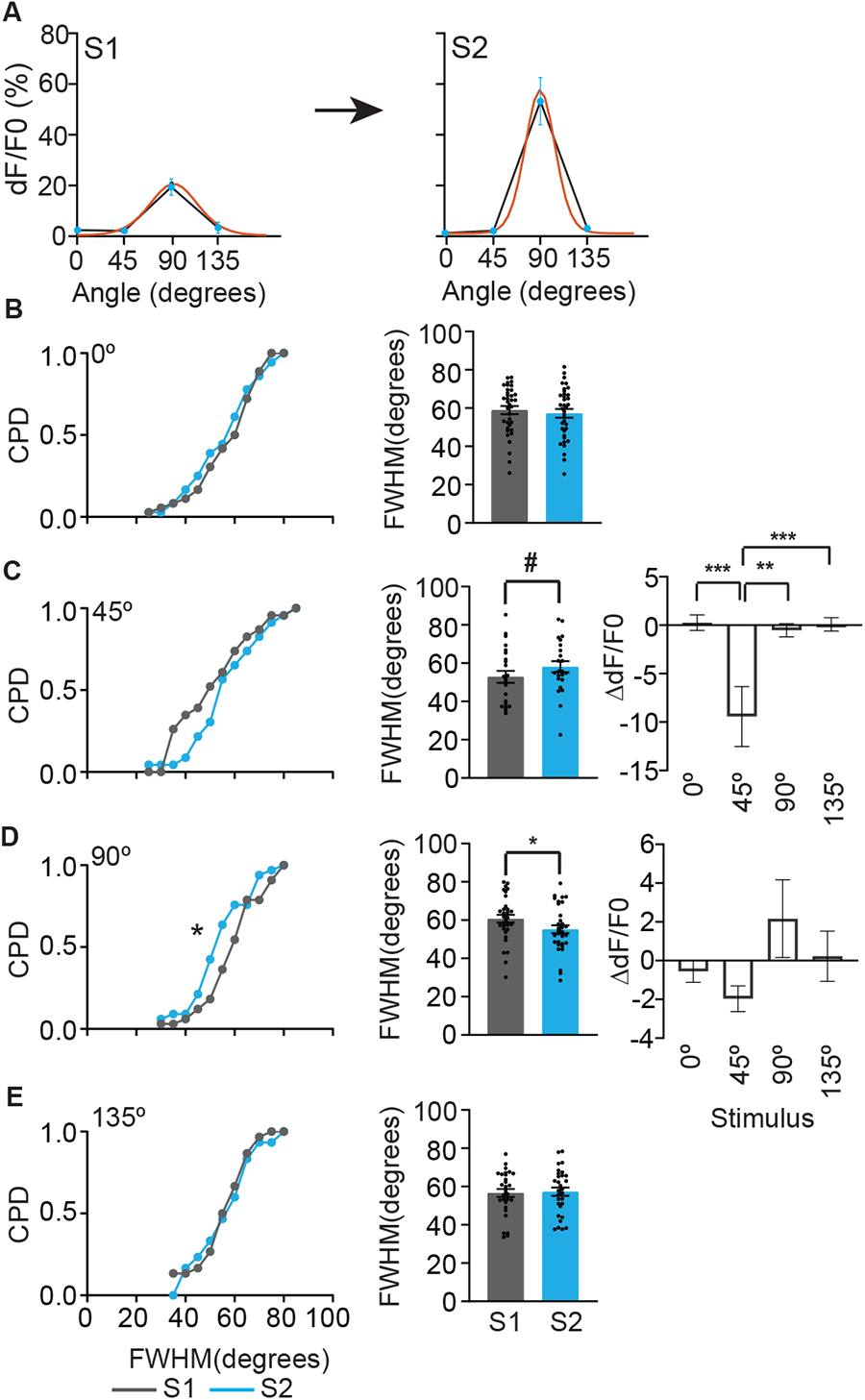
Altered experience elicits stimulus competition in high-responsive neurons. **A.** Representative orientation tuning curve fit (orange line) of a highly responsive neuron whose tuning curve sharpened following altered experience. The blue circles represent the mean of trial-averaged dF/F0 of 0°, 45°, 90°, and 135° grating stimuli, and the error bars represent SEM. **B-E.** Left. Cumulative probability distribution (CPD) of full width at half maximum (FWHM) of matched neurons that retained high responsiveness and preferred orientation before (S1) and after (S2) altered experience. * *p*<0.05, KS test; *n* = 36, 23, 33, and 30 matched neurons that remained highly responsive and tuned to 0°, 45°, 90°, and 135°, respectively. Middle. Mean FWHM of the same neurons. # *p* <0.06, * *p* < 0.05, linear mixed effects model. Black circles represent individual neuron values. (C and D) Right. Change in mean trial-averaged dF/F0 (ΔdF/F0) following altered experience for the neurons tuned to 45° (C) or 90° (D). * *p* < 0.05, ** *p* < 0.01, ** *p* < 0.01, *** *p* < 0.001, linear mixed effects model followed by Tukey’s post hoc multiple comparisons test. Data are presented as mean ± SEM.

To test whether the sharpening and broadening of the tuning curve is a result of stimulus competition, we compared the average ΔdF/F0 of these neurons. We found that the broadening of the tuning curve of familiar stimulus-preferring neurons is solely due to a reduction in familiar stimulus responsiveness. In contrast, the sharpening of the tuning curve of 90° preferring neurons is due to a subtle increase in 90° response and a slight reduction in 45° response, indicative of stimulus competition (Fig. 3C, D). These results show that competition in highly responsive neurons primarily occurs in neurons tuned to orientations up to 45° apart from the familiar orientation, though it did not reach significance for 0°-tuned neurons.

### Neurons tuned to orientations 45° apart from the familiar stimulus dominate functional connectivity

To assess whether stimulus competition elicited by altered experience changes the functional connectivity of neurons, we identified neurons with significant coactivity during the entire imaging period consisting of multiple visual stimuli (Fig. 4A). Two neurons are functionally connected if the number of their coactive imaging frames (imaging frames with deconvolved spikes for the compared neuron pair) is >95% of the cumulative distribution of their coactivity obtained by 1000 random circular shifts of the imaging frames^48^. For each session, we calculated the functional connectivity (node degrees) based on the tuning preference of neurons in that session. We did not see a significant change in the average node degrees for neurons tuned to any orientation, including those tuned to familiar orientation or all combined (Fig. 4B). This was surprising because we previously found a reduction in node degrees following familiarity^4^. However, in that study, functional connectivity was assessed for all the neurons within the familiar stimulus period, whereas our current analysis encompasses imaging frames spanning multiple stimuli. Thus, we could detect functional connectivity associated with non-familiar and familiar stimuli. To confirm this, we limited our analysis to imaging frames corresponding to familiar stimulus experience and found that functional connectivity was reduced following familiarity (Fig. 4B (bottom right)).

**Figure 4.**
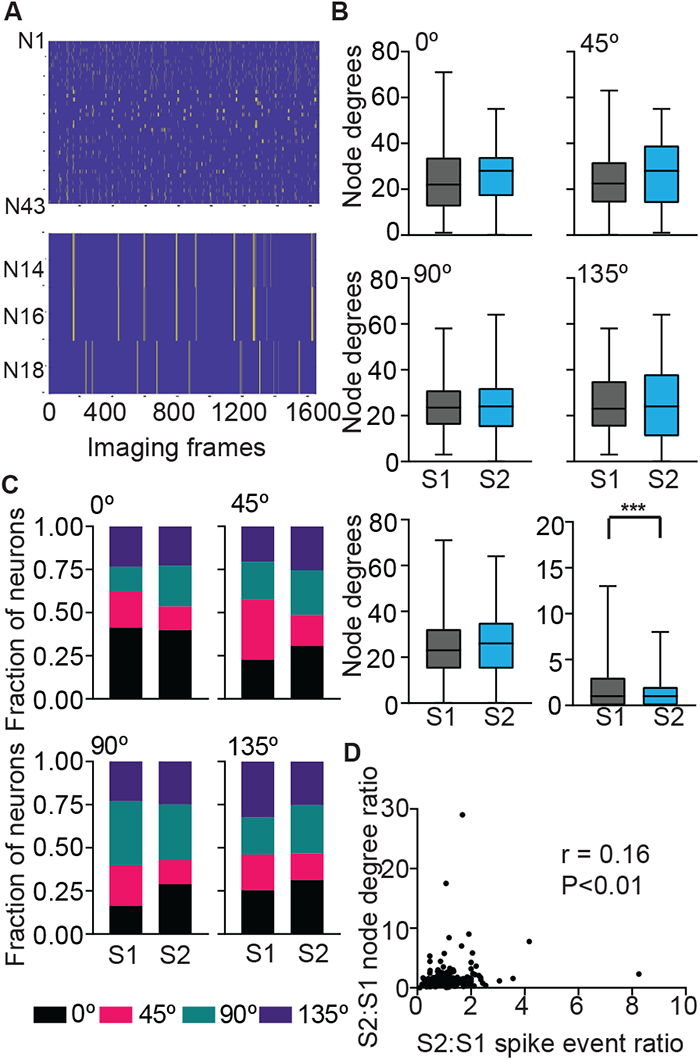
Neurons tuned to non-familiar stimuli dominate functional connectivity. **A.** Representative raster plot of binarized deconvolved spikes from 43 neurons (N1-N43) neurons of one mouse. A plot of three of those neurons (N14,16,18) is magnified below. N14 and N16 are functionally connected to each other but not N18. **B.** Average node degrees across the entire imaging period of active neurons with good curve fit tuned to 0°, 45° (top), 90°, 135° (middle), combined (bottom left) before (S1) and after (S2) altered experience. Average node degree of the same combined neurons when restricted to the 45° stimulus period. Data are presented as box (25th to 75th percentile) and whisker (minimum and maximum values) plots, the median value indicated as a horizontal line. *n* = 10 mice (104, 123 (0°), 96, 52 (45°), 94, 109 (90°), 88, 95 (135°), 382,379 (combined) neurons (S1,S2). *** *p* < 0.001, linear mixed effects model. **C.** The fraction of partner neurons with different tuning preferences for seed neurons tuned to the indicated orientation. **D.** Correlation between the ratio of deconvolved spikes and node degrees between sessions for 274 matched neurons that had at least one functionally connected neuron in both sessions.

To test whether functional connectivity is reorganized despite no change in average functional connectivity across the entire imaging session, we first identified the tuning preferences of the functionally connected partner neurons for each neuron (seed neuron). We grouped the seed neurons based on their tuning preference and calculated the distribution of tuning preferences of their partner neurons. We found that ∼30-40% of partner neurons share the same tuning preference as the seed neurons. After the altered experience, the fraction of partner neurons tuned to familiar orientation decreased for seed neurons tuned to any orientation. Interestingly, the fraction of partner neurons tuned to 0° and 90° increased by ∼70% for seed neurons tuned to 90° and 0°, respectively, following altered experience (Fig. 4C). Similarly, seed neurons tuned to 45° and 135° showed ∼20-30% increase in partner neurons tuned to 0° or 90° (Fig. 4C). Changes in functional connectivity could reflect reorganization of firing patterns or could be a reflection of altered firing rate. We found that the change in the number of deconvolved spikes correlated weakly (but significantly) with the change in functional connectivity for neurons matched across sessions (Fig. 4D), indicating that altered responsiveness may not solely be responsible for robust functional connectivity reorganization. These results indicate that stimulus competition associated with familiarity alters functional connectivity favoring the dominance of neurons tuned to stimuli that are 45° apart from the familiar stimulus.

### Responsiveness to natural images is altered following familiarity with a grating stimulus

Changes to orientation grating stimulus responsiveness could influence neural responsiveness to natural images, which consist of multiple orientations. To test this possibility, we compared the fraction of neurons responsive to natural images before and after the altered experience with the 45° stimulus. We found that the fraction of non-responsive neurons reduced and transitioned to low responsiveness to the set of ten natural images tested (Fig. 5A-C). Interestingly, the neurons that became responsive (gain) to these natural images also showed increased responsiveness to non-familiar grating stimuli, consisting of cardinal orientations, and decreased responsiveness to the familiar grating stimulus. In contrast, neurons that became non-responsive (loss) after altered experience or those that remained stably responsive had no or minor changes to non-familiar stimuli responsiveness (Fig. 5D). These results suggest that an increase in responsiveness to 0° and 90°, presumably due to stimulus competition associated with familiarity, alters responsiveness to natural image stimuli.

**Figure 5.**
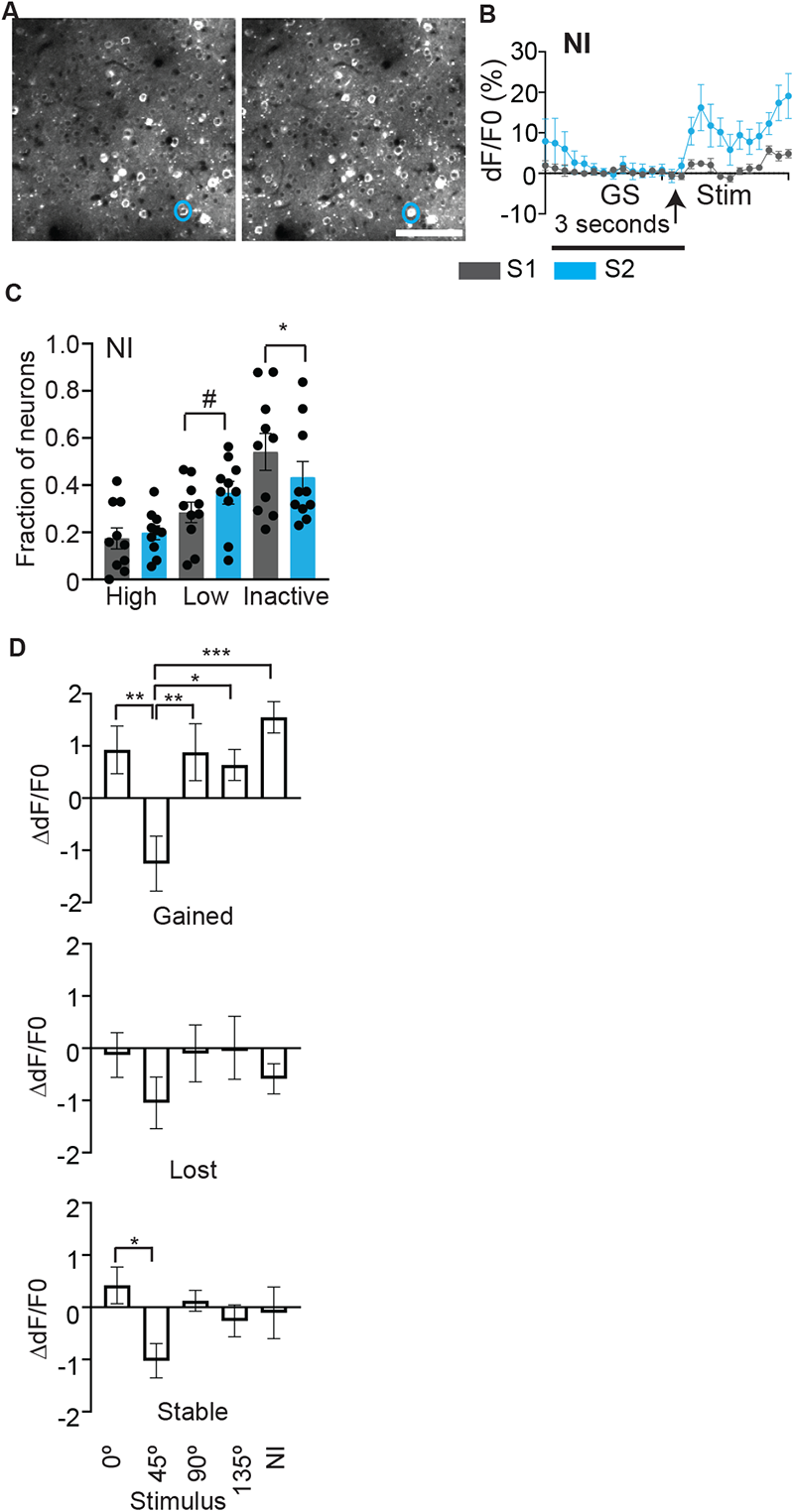
Increased fraction of neurons responsive to natural images following altered experience. **A.** Representative imaging fields on S1 (left) and S2 (right) from one mouse. Scale bar: 100 µm. **B.** Trial-averaged dF/F0 ± SEM elicited by natural images from the neuron circled in blue in A. **C.** Fraction of high, low, and non-responsive neurons to natural images. Circles on the histogram represent individual mouse values. *n* = 10 mice. # *p* = 0.05, * *p* < 0.05, paired *t* tests. **D.** Change in dF/F0 (S2 – S1 dF/F0) in response to 0, 45, 90, and 135° stimuli in neurons that became responsive (gain), unresponsive (loss), or remained active (stable) to natural images. * *p* < 0.05, ** *p* < 0.01, *** *p* < 0.001, linear mixed effects model followed by Tukey’s post hoc multiple comparisons test. Data are presented as mean ± SEM.

### Spontaneous activity patterns reflect the altered experience

We next compared whether the spontaneous activity is modulated by altered experience in neurons that retained high responsiveness to stimuli (Fig. 6A). For this analysis, imaged neurons were matched between the spontaneous period and evoked activity images independently for pre- and post-altered experience imaging sessions. Spontaneous activity was quantified as the average dF/F0 obtained from every 3 seconds bin (to match the average stimulus duration) over the ∼237 seconds of imaging in darkness. We found that the average dF/F0 of spontaneous activity did not change in familiar or non-familiar preferring neurons following the altered experience, and the response between sessions was not correlated (Fig. 6B). Though the average magnitude of spontaneous activity may not be significantly different, the relative activity levels of individual neurons in the population could change following an altered experience. We calculated the similarity ratio to assess changes to cosine similarity (described in Methods) between spontaneous and stimulus-evoked activity in highly responsive neurons. An increase in the ratio above one indicates increased similarity between spontaneous and evoked activity following altered experience. The similarity between spontaneous and 0°, 45°, 90°, 135°, and gray screen evoked responses were 0.45, 0.27, 0.31, 0.34, and 0.38, respectively, on S1. It changed to 0.43, 0.38, 0.27, 0.36, and 0.55 on S2 for the same stimuli (Fig. 6C). When restricted this analysis to neurons based on their tuning preferences, the similarity of spontaneous and preferred stimulus-evoked activity increased by three-fold for familiar stimulus preferring neurons (0.27 to 0.84 similarity). Spontaneous activity and the preferred-stimulus evoked response were greater for horizontal orientation before the altered experience (0°; 0.63 similarity) but moderately reduced to 45% following the altered experience. The similarity of spontaneous activity and preferred stimulus-evoked response in neurons tuned to 90° (0.38 on S1) and 135° (0.36 on S1) stayed approximately the same (Fig. 6D).

**Figure 6.**
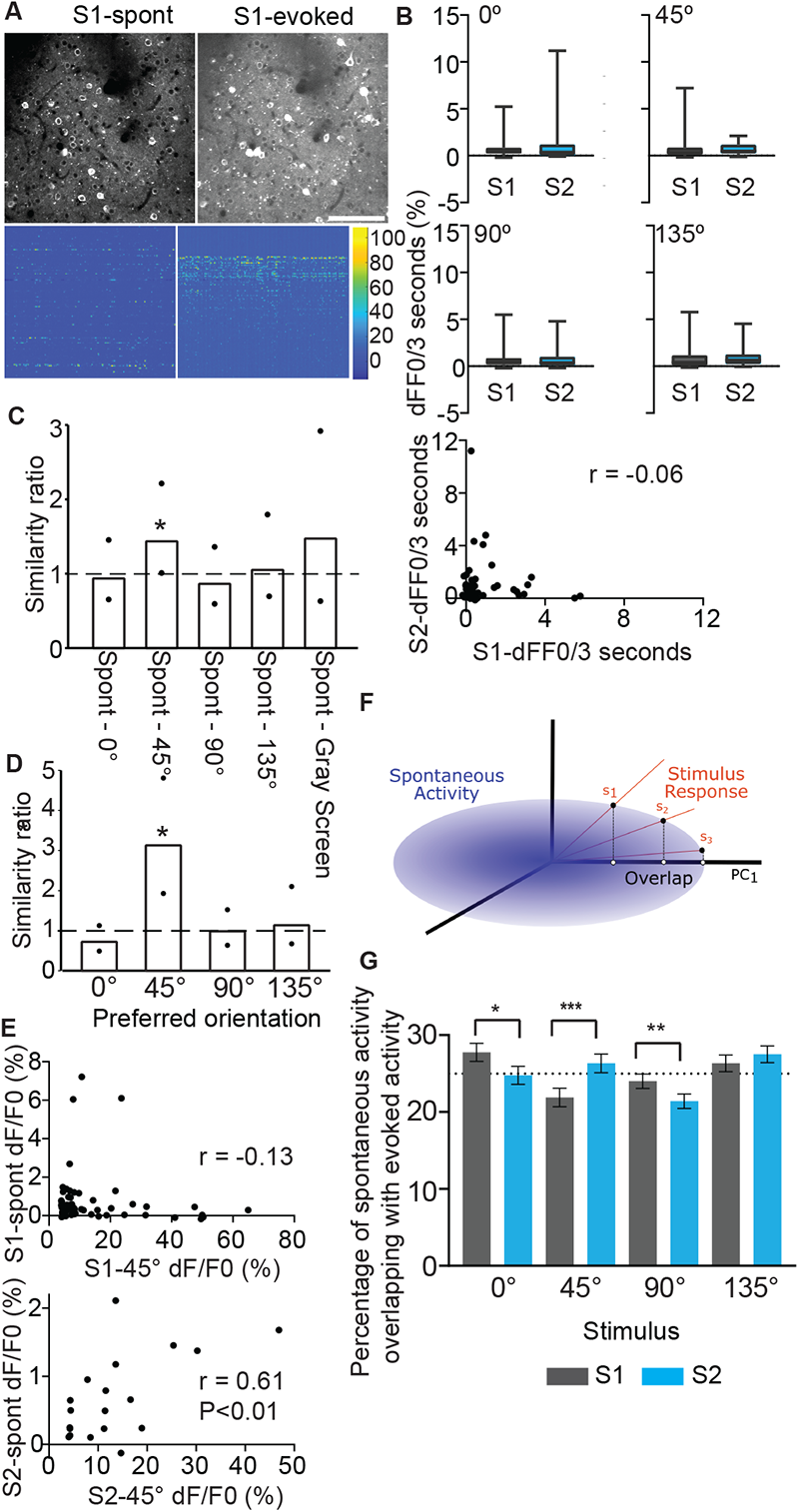
Spontaneous and familiar stimulus-driven activity patterns overlap. **A.** A representative standard deviation projection images from spontaneous and evoked imaging sessions (top; Scale bar: 100 µm) and corresponding dF/F0 (color bar – dF/F0) raster plot (below). **B**. dF/F0 averaged over three seconds during spontaneous activity in highly responsive neurons tuned to the indicated orientation before (S1) and after the altered experience (S2). Data are presented as box (25th to 75th percentile) and whisker (minimum and maximum values) plots, with the median value indicated as a horizontal line. n = 50, 53, 50, 38 (S1) and 43, 19, 34, 40 (S2) neurons tuned to 0°, 45° (top), 90°, 135° (middle) bins, respectively from 9 mice. Bottom – Correlation of average dF/F0 of spontaneous activity of highly responsive neurons matched for spontaneous and evoked imaging in both S1 and S2 (53 neurons). **C.** Fold-change in the similarity of responses to spontaneous activity and evoked activity elicited by indicated grating stimulus or gray screen (GS) in all neurons highly responsive to any grating stimulus. *n* = 191 (S1) and 136 (S2) neurons from 9 mice. **D.** Fold change in the similarity of responses to spontaneous and evoked activity elicited by indicated stimulus in neurons tuned to that stimulus. *n* = 50, 53, 50, and 38 neurons (S1) and 43, 19, 34, and 40 neurons (S2) tuned to 0°, 45°, 90°, and 135°, respectively from 9 mice. The dotted line indicates identical similarity after 45° experience. Black circles – the lower and upper bound of 95% confidence intervals (CI) obtained by bootstrapping with replacement. * null value (1.00) outside of 95% confidence interval. **E.** Correlation between average 45° evoked dF/F0 and average spontaneous dF/F0 for highly responsive neurons tuned to 45° (n = 53 neurons (S1), 19 neurons (S2). F.Schematic for calculating the overlap between evoked responses to stimuli and spontaneous neural activity. The spontaneous neural activity is defined by the covariance matrix represented by the blue ellipsoid. Each trial of stimuli can then be represented as a vector defined as the response activity across all observed neurons. The calculated overlap between these stimuli responses and the spontaneous activity was normalized relative to the 1st principal component with maximum variance. **F.** Percentage of spontaneous neural activity that overlapped with the evoked responses for the four grating stimuli. The overlap for each of the stimuli for each time window was divided by the total overlap observed across all four stimuli in a cycle to show the relative percentages of spontaneous activity that mimicked each of the stimuli. The dotted line indicates an equal amount of overlap (25%) across the four stimuli. After the altered experience, neural activity during the spontaneous period significantly increased its overlap from S1 to S2 to that evoked with the familiar 45° stimulus and decreased for the 0° and 90° stimuli (*p<0.05, **p<0.01, ***p<0.001 two-way ANOVA, posthoc Tukey test). Data are presented as mean ± 95% confidence interval from the ANOVA analysis of eight repeated stimuli cycles for nine mice x two sessions.

To test whether the reduction in responsiveness to the familiar stimulus alone contributes to increased cosine similarity, we restricted our analysis to neurons present in spontaneous and evoked imaging sessions in S1 and S2 and that were highly responsive with good tuning curve fit. These neurons (tuned for any of the grating stimuli; n = 53 neurons) also show increased cosine similarity between spontaneous and familiar stimulus-evoked activity (0.14 (S1) and 0.37 (S2) similarity). Importantly, the mean response to 45° stimulus (mean trial-averaged dF/F0 (%): 8.7±2 (S1) and 6.4±1.2 (S2)) and average spontaneous activity (dF/F0 (%) per 3 seconds: 0.29±0.8 (S1) to 0.58±1.03 (S2)) were not significantly different, indicating increased similarity is due to reorganization of activity rather than changes to average activity. To further demonstrate the reorganization of activity relationships, we chose highly responsive neurons tuned to 45° stimulus and performed a correlation analysis between their average response to 45° stimulus and spontaneous activity. We found that repeated experience of 45° stimulus resulted in an enhanced correlation between familiar stimulus-evoked and spontaneous activity, indicating activity reorganization in the population (Fig. 6E).

We also measured the stimulus-evoked and spontaneous activity overlap for individual three-second time bins. We projected the neural response of stimulus-evoked activity for individual stimulus cycles onto the observed spontaneous activity neural space (described in Methods; Fig. 6F). The overlap of spontaneous activity with each of the four grating stimuli evoked responses (obtained from all responsive neurons) was evenly distributed with a slight propensity for 0° on the first day of imaging. After the altered experience, however, the spontaneous neural activity increased its overlap with that of the familiar 45° stimulus with a corresponding decrease in overlap with the 0° and 90° stimuli (Fig. 6G). In the cartoon in Fig. 6F, the overlap between the vector and ellipse is changed only by relative changes between activity levels of neurons. When all the neurons’ activity changes by the same amount, the ellipse would rescale; however, both the vector and ellipse are normalized to total activity, so they are always a unit vector or unit volume. Therefore, the overall size never changes in the overlap calculation, even when the global activity changes. Thus, the increase in overlap indicates a reorganization of neural activity patterns rather than an average global reduction in activity. Thus, neural activity patterns that occurs when no stimuli are present and when the familiar stimulus was experienced more closely mimicked each other.

## Discussion

Consistent with a vast body of literature on habituation, we show that repeated orientation-grating stimulus experience reduces neural responsiveness to the familiar stimulus in the mouse visual cortex. Neurons in the visual cortex receive functionally diverse inputs, with neighboring synapses tuned to different orientations^16,19^. Long-term reduction in responsiveness to one set of inputs evoked by familiarity is likely to elicit a compensatory increase in other inputs, perhaps to maintain neuronal activity homeostasis. Competition between stimuli is well established using different paradigms, such as monocular deprivation^49,50^, cross-modal plasticity ^51-54^, and sensory adaptation^20,25,26^.

In contrast, there is wide variability in reported findings on how multi-day repeated passive visual experience influences neuronal activity and tuning^3-5,8,11-13,15^. Even among studies that used calcium imaging as a proxy for neural activity, familiarity is associated with an increase^12^, decrease^4,13-15^, or no change^11^ in neural activity. The use of anesthesia^12^, the effects of locomotion, and other variables associated with the stimulus type, imaging, and data analyses could contribute to these differences. The majority of these studies, however, found a reduction in responsiveness to the familiar stimulus, and the current study adds more support to these observations. However, the studies that found reduced responsiveness to the familiar stimulus did not report altered tuning properties. Consistent with these studies, we also did not observe a significant shift in tuning preference when we analyzed all neurons considered to be visually responsive or when limited to highly responsive neurons. We only observed a significant shift in tuning preference when we restricted our analyses to neurons that were low responsive before and after the altered experience. This is consistent with the proposal that plasticity may differ based on activity levels during passive viewing^55,56^.

The greater vulnerability of lower responsive neurons to familiarity-evoked stimulus competition could be because weaker synapses are more likely to undergo potentiation or depression^57,58^. Alternatively, subtle changes to inhibition that suppress inputs tuned to familiar orientations and disinhibit inputs tuned to nearby orientations could have a higher impact on lower responsive neurons. Depression and potentiation of familiar orientation and nearby non-familiar orientation stimulus-responsive synapses in lower responsive neurons could lead to fewer retaining the familiar orientation as the preferred stimulus. Similar plasticity mechanisms, albeit to a lesser extent, in highly responsive neurons may underlie subtle sharpening of tuning width of neurons tuned to 90° stimuli. Due to the slightly lower sampling of orientation tuning space in this study, we cannot rule out that the increased tuning width is due to a subtle shift in tuning preference.

Neural responsiveness, the fraction of neurons tuned to different orientations, tuning width, and functional connectivity were strongly reduced, whereas similarity with spontaneous activity was selectively increased in neurons tuned to 45° stimulus compared to other stimuli, suggesting repeated stimulus experience rather than time is likely a major contributor for the observed changes. The subtle increase in the responsiveness of neurons tuned to 90° and the dominance of neurons tuned to 0° or 90° in forming functional connections following altered experience could be due to their relative proximity to the experienced stimulus compared to 135° stimulus. Alternatively, the increased dominance of 0° and 90° tuned neurons could be due to the enrichment of these angles in the animal’s visual environment and, therefore, are ethologically more relevant. Whether the repeated experience of cardinal angles would lead to a dominance of ethologically less enriched orientations remains to be tested.

What purpose could stimulus competition serve? Habituation allows resources to be spent on detecting novel or behaviorally relevant stimuli. Models of habituation that involves synaptic depression or enhanced inhibition could explain a reduction in familiar stimulus responsiveness but do not explain how novel stimuli are preferentially detected and how a reduction in neural activity generates the knowledge of familiarity^1,59^. Predictive coding-based models suggest that the internal model of visual experience allows for the selective transmission of sensory information that deviates from the expectation^60-63^. The internal model could serve as the knowledge of familiarity, and their inhibitory influence on familiar stimulus responsiveness may contribute to novelty detection. Consistently, deviation from or memory of expected sensory experience has been shown to modulate neuronal activity in the visual cortex^30,38,61,64-71^. Similarly, spontaneous activity also influences stimulus-evoked activity^72,73^. In contrast, our findings indicate that the internal model of familiarity and enhancement of novel stimuli responsiveness may occur independently. We found that spontaneous and stimulus-evoked activity structures become more similar if the stimulus is familiar. The increase in similarity is more prominent in familiar stimulus-preferring neurons. These results suggest that the internal model or expectation of familiarity may arise when spontaneous and familiar stimulus-evoked activity patterns are similar. We suggest that the suppression of familiar stimulus responsiveness occurs independent of changes in spontaneous activity. We show that the average amplitude of spontaneous activity is not altered by familiarity, and the change in the similarity between the prestimulus gray screen elicited and spontaneous activity is not significant. The amplitude of familiar stimulus-evoked activity, much larger than that of prestimulus gray screen activity, is reduced. Instead, we found that suppression and enhancement of familiar orientation and nearby non-familiar orientation stimuli responsiveness go hand-in-hand, presumably driven by stimulus competition. However, further experiments are needed to establish the independence of spontaneous and evoked activity changes following familiarity. Our observations contrast with a recent study that showed increased spontaneous but no change in evoked activity in the visual cortex following multiple days of grating stimulus exposure in head-fixed but not body-restrained mice. Interestingly, the spontaneous activity and behavioral habituation increase did not remain specific to the experienced orientation^11^. It is unclear whether the differences in the results are due to locomotion, which is known to influence neuronal activity in the visual cortex significantly^74^, stimulus features (drifting vs. phase reversing gratings), or body restraint-induced stress-modulated cortical activity.

The increased responsiveness and higher functional connectivity of neurons tuned to some non-familiar stimuli reflect the detection of deviation from the expected familiar stimulus. In this view, error detection is built in the neurons by stimulus competition and occurs independently of the spontaneous activity representation of the expected or familiar stimulus (Fig. 7). Though not directly addressed in this study, we also speculate that stimulus competition could explain neural activity that persists when an expected stimulus is skipped^65,75,76^ if the direction of competition between familiar and non-familiar inputs is opposite in excitatory and specific class of inhibitory neurons (Fig. 7). This speculation is based on studies that show stronger familiar stimulus-evoked activity in somatostatin expressing inhibitory neurons^77^, which are also involved in detecting skipped expected stimulus^78^. In this model (Fig. 7), reduced excitation and increased inhibition by the familiar stimulus will weaken neural activity, whereas increased excitation and reduced inhibition by novel stimuli will enhance it. Furthermore, skipping an expected stimulus will lead to disinhibition and elicit neural activity.

**Figure 7.**
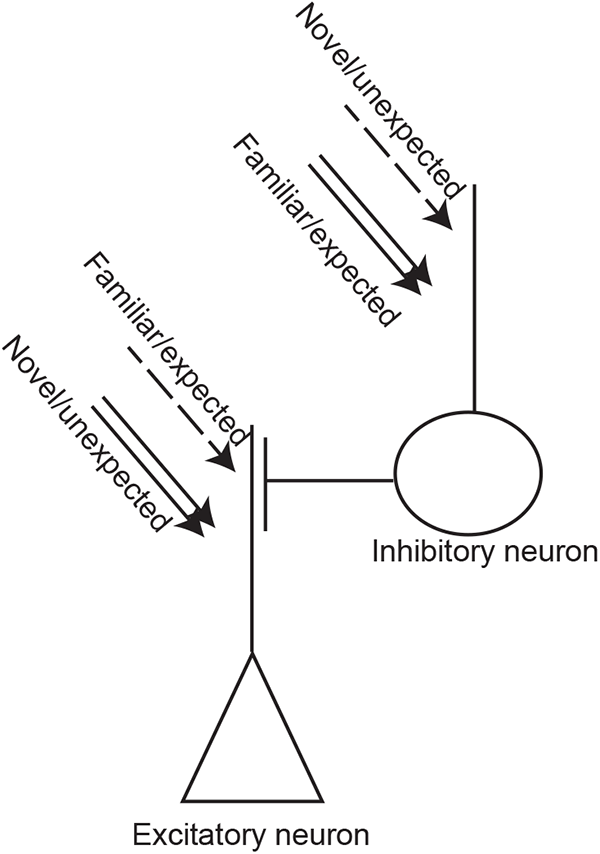
A model for error detection enabled by stimulus competition. Repeated passive experience may weaken familiar or expected stimulus inputs and strengthen novel or unexpected stimuli inputs onto excitatory neurons. However, the direction of this change could be the opposite in certain classes of inhibitory neurons. Reduced excitation and enhanced inhibition elicited by familiar or expected stimulus will dampen neuronal activity; skipping the expected stimulus will lead to disinhibition and restore activity. In contrast, novel or unexpected stimuli will elicit higher neuronal activity. Double arrows and dashed arrows indicate the strengthening and weakening of synaptic inputs following passive exposure, respectively.

Beyond enabling novelty detection, stimulus competition may contribute to memory generalization or unintended learning. We found a subtle increase in the fraction of neurons responding to the same natural images after familiarity with a 45° grating stimulus. Natural images consist of multiple orientations with a predominance of cardinal angles^79^, whose responsiveness was also increased in neurons activated by natural images post familiarity. We propose that non-learned stimuli with features that compete with or are similar to the familiar stimulus will increase and decrease neuronal responsiveness.

## Methods

### Mice

All animal procedures are approved by the University of Kansas Institute of Animal Use and Care Committee and meet the NIH guidelines for the use and care of vertebrate animals. C57BL/6J-Tg (Thy1-GCaMP6s) GP4.3Dkim/J (https://www.jax.org/strain/024275) were maintained as heterozygotes ^80^. A maximum of five mice were housed in a standard cage but individually housed after the cranial window surgery. Mice were housed on a 12h-light/12h-dark cycle. Isoflurane was used as the anesthetic for cranial window surgeries, and C0_2_ was used for euthanasia.

### Cranial window

∼ 4-month-old GCaMP6S mice received a cranial window on the right hemisphere over the visual cortex. A small scalp incision was made over the midline of the skull. A 5-mm diameter circle covering the visual cortex was scored using a biopsy punch. The skull was thinned along the scored circle with a fine drill using a sterile 0.5mm diameter round burr (Fine Science Tools). The bone flap was removed with fine forceps leaving behind the dura. A 5-mm diameter sterile circular glass coverslip (Harvard Apparatus) was positioned over the opening. Vetbond and cyanoacrylate glue was applied between the coverslip and bone to keep the coverslips in place. Metabond (C&B Metabond) was applied over the exposed skull. ∼2-weeks after the surgery, a titanium head-post was affixed around the window to restrain mice during imaging. A black paint powder was applied over the cement, and a light-blocking cone was attached to the titanium headpost to block monitor light from reaching the photomultiplier tubes (PMTs) during imaging of visually evoked activity.

### Widefield calcium imaging

Widefield calcium imaging was performed 14 days after cranial window surgeries to identify the visual cortex. Imaging was performed in a custom-built upright microscope with a 4X objective (Nikon). Awake mice, habituated for two days to the microscope, were positioned 20 cm in front of a high refresh rate monitor displaying a horizontal bar (1° of the visual field) drifting at 10 Hz. Images were collected using an sCMOS camera at 5Hz (1024 x 1024 pixels; Photometrics). GCaMP6 was excited by an LED (Lambda FLED, Sutter) filtered through a bandpass filter (470/40, 49002 Chroma), and the emission was filtered through a 525/50 bandpass filter. Reference vasculature was imaged with a 470 nm fiber-coupled LED powered by T-Cube LED drivers (Thorlabs). Images were downsized to 256×256 pixels, and magnitude maps, based on GCaMP6 fluorescence, were computed by extracting the Fourier component of fluorescence changes to matched stimulus frequency. The fractional change in fluorescence represents response magnitude, and the magnitude maps were thresholded at 30% of the peak-response amplitude. The visually responsive cortex was mapped by overlaying the magnitude maps over the 470nm reference image.

### Two-photon imaging

Neurons within the mapped visual cortex (∼100-150 μm below the dura) were imaged at 4.22 Hz, using a Sutter MOM multiphoton microscope, in head-fixed awake mice restrained in a body tube. The Ti: sapphire laser (MaiTai HP: Newport SpectraPhysics; 940 nm) was routed to the microscope using table optics. The power was adjusted (20-40mW) using a rotating half-wave plate and a polarizing beam splitter to avoid signal saturation. A pair of galvanometric mirrors scan the laser beams to the back aperture of the objective (Nikon 16X 0.8 NA). The emission signal was collected through the same objective, passed through a short pass filter to block infrared wavelengths, and routed to a GaASP PMT after passing through a 540/50 bandpass filter. Image acquisition was controlled by Scanimage (Vidrio Technologies). The imaging field was a single Z frame of 336 x 336 um (256 x 256 pixels) consisting of 100 or more cells.

### Visual stimulus to head restrained mice

Visual stimuli were delivered on a high refresh rate monitor placed 20 cm in front of the head-restrained animals covering 94° x 61° of the visual field. The software for generating visual stimuli was modified from a custom-written stimulus suite (a kind gift from Dr. Mark Bear’s lab) written in Matlab (Mathworks) using the PsychToolbox extension (http://psychtoolbox.org). Mice were habituated to a gray screen by head-restraining them under the microscope for two days (30 minutes each day). On the first imaging session (S1), 10-15 minutes after head fixation, the GCaMP6 response was first imaged without visual stimulus (total darkness) to record spontaneous activity. ∼2 minutes later, visually evoked activity imaging was performed. Visual stimuli consisted of 30 seconds of the gray screen followed by 8-repetitions of 100% contrast, sinusoidal, phase reversing (2 Hz, 0.05 cycles /degree) grating stimuli of different orientations (0°, 45°, 90°, 135° – 3 seconds each) and a set of ten natural images (0.3 second/image – 3 seconds per set) interspersed with 6 seconds of the gray screen. The order of stimuli was different in each cycle. Grayscale natural images were obtained from Berkeley Segmentation Dataset, contrast normalized, and resized to 1600 x 1068 pixels. Grating stimuli covered the entire monitor display value range between black and white. Gamma correction was performed to ensure the total luminance in the gray screen and grating stimuli were the same.

For the next eight days, head-restrained mice were exposed to two sessions of 60 seconds of the gray screen followed by five blocks of 100 seconds of phase-reversing grating stimulus with 30 seconds of the gray screen between blocks. The two sessions were separated by ∼1-2 hours. On the following day (S2), spontaneous and evoked activity was imaged as before. We tried to closely match the same field of view imaged in the first session for post-training imaging on the ninth day (from S1).

### Calcium imaging analysis

Motion registration and ROI detection in the time-series images were performed using Suite2p RRID: SCR_016434 ^46^. Tau and neuropil coefficient for spike deconvolution were set at 2.0 and 0.5, respectively. Suite2p generated ROIs were chosen as cells (cellular ROI) if the soma was visible in the mean or maximum projection image. All suite2p ROIs were matched between sessions. The following imaging sessions were matched for the analyses – S1-evoked and S2-evoked; S1-spontaneous and S1-evoked; S2-spontaneous and S2-evoked. Spontaneous and evoked sessions were separated by 5-10 minutes and therefore were matched to ensure no Z movement occurred during this time. If an ROI is present in one of the matching sessions but not the other, then a manual ROI was placed if the mean projection image showed the presence of a morphologically similar neuron at the same location.

Cellular fluorescence (F) was corrected for neuropil contamination, estimated as the ratio of blood vessel fluorescence to that of neuropil (Fneu). Neuropil-corrected fluorescence (Fcorr) was calculated as F – (0.5xFneu). dF/F0 is calculated as (Fcorr – F0)/F0, where F0 is defined as the mode of the Fcorr density distribution across the entire contiguous imaging session. F0 is calculated independently for spontaneous and evoked imaging sessions. Cellular ROIs that did not have at least one peak greater than 10% dF/F0 anywhere in the time series in at least one of the matched sessions were excluded. The 10% dF/F0 could lie anywhere in the time series (corresponding to the gray screen or the stimulus period) and is averaged out by the trial variation. We then manually examined all matched neurons and only selected ones with clearly visible soma in both sessions.

For each neuron, dF/F0 elicited by each stimulus was calculated as the mean dF/F0 of imaging frames corresponding to three seconds of the stimulus period and the preceding three seconds of the gray screen from the eight trials. The mean of trial-averaged dF/F0 of a neuron for each stimulus and the preceding gray screen was calculated as the mean of average dF/F0 during the three-second stimulus and three-second gray screen periods, respectively. Neurons are considered active if the mean of trial-averaged dF/F0 during the three-second (13 imaging frames) stimulus period is greater than two standard deviations of the mean of the trial-averaged dF/F0 of the three-second (13 imaging frames) gray screen preceding it (p<0.05, paired t-test to compare 13 imaging frames of the gray screen and stimulus periods). Neurons are considered high or low responsive if they pass the criteria for active neurons and have a trial-averaged mean dF/F0 of >4 or <4, respectively, for at least one of the grating stimuli.

To obtain an orientation tuning curve, the trial-averaged area under the curve of dF/F0 during the 3 seconds of each grating stimulus was fit as a function of stimulus angle (L) with a von Mises function (Eqn. 1) in neurons responsive to any grating stimuli.

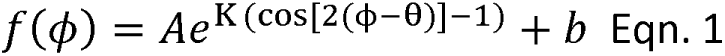

The function is defined by four fit parameters: a preferred stimulus orientation that gives the maximum response (θ), a tuning curve width (Κ), a response amplitude (A), and an intercept (b). Note, in our equation the angle difference (☐ – θ) is doubled to fit the 180☐ data to the standard 360☐ von Mises function. Fits were calculated with a maximum likelihood estimate of θ and Κ of using CircStat in Matlab and least-squares regression was then used to identify A and b. The fraction of explained variance, R^2^, was calculated, and the full width at half-maximum (FWHM) was calculated as:

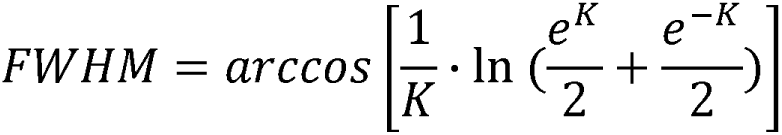

Note, this equation is one half of the standard equation for FWHM of a von Mises function due to our range of angles from 0☐ to 180☐. The amplitude (Amax, Eqn. 2) at the preferred stimulus angle (θ) and amplitude (Amin, Eqn. 3) at the opposite angle, 90☐ π/2 radians from preferred, were also calculated.

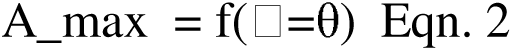

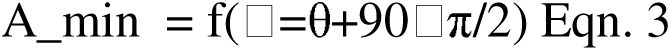

Only neurons with a tuning curve fit with R^2^>0.7 and were identified as visually responsive were used for analyses involving preferred orientations.

The fraction of neurons preferring an indicated orientation is calculated as the number of neurons whose preferred orientation is within 22.5° of the indicated bin divided by the number of neurons in all the bins. To calculate the fraction of neurons that retained the original or changed their preferred orientation, we binned (± 22.5° from the indicated orientation) neurons based on their S1 preferred orientation and calculated the number of matched neurons on S2 with the indicated preferred orientation divided by the total number of neurons in that group. To binarize as 45° and non-45° neurons, we pooled neurons tuned within 22.5° to 67.5° as 45° preferred neurons and the rest as non-45° neurons. The fraction of neurons responsive (active) for natural images was calculated as the number of neurons considered active for natural images divided by the total number of identified neurons.

Paired FWHM of matched highly responsive neurons on S1 and S2 evoked sessions was calculated as the mean FWHM of all matched highly responsive neurons that had the preferred orientation within the same bin on both sessions. ΔdF/F0 was calculated as mean trial averaged dF/F0 on S2 – S1 for the same stimulus for each neuron.

Deconvolved spikes obtained from Suite2p for cellular ROIs were thresholded (> 2 SD from the mean) and binarized to assess functional connectivity. Briefly, neuron pairs are considered functionally connected if the number of their coactive frames exceeds 95% of the cumulative probability distribution generated by a 1000 random circular shift of their activity. The functional connectivity matrix generated above was used to determine the node degree – the number of edges connected to each node (neuron) during the entire imaging period using MATLAB graph and degree functions. To assess functional connectivity selectively during the 45° stimulus period, we only kept the frames corresponding to that stimulus.

To quantify spontaneous activity, the mean dF/F0 for each three seconds imaging period in darkness was first calculated for each neuron. The mean of 39 dF/F0 was obtained for each neuron. Neurons were matched with the evoked session on the same day to classify them in preferred orientation bins. Mean spontaneous dF/F0 was calculated for matched high responsive neurons tuned to the indicated orientation in the corresponding evoked session.

The cosine similarity between neural responsiveness to spontaneous and the indicated stimulus was calculated as similarity = X.Y/||X||||Y||, where X and Y are the average spontaneous dF/F0 and the mean of trial-averaged dF/F0 elicited by the indicated stimulus in highly responsive neurons. The same analysis was restricted to highly responsive neurons tuned to the indicated bin orientation to identify the similarity of responses based on tuning preference. Neurons matched between spontaneous and evoked imaging on S1 and S2 were used for this analysis.

We also measured how much spontaneous neural activity overlapped with the stimulus-driven response from the grating patterns for individual three-second time windows. We defined the neural subspace of spontaneous activity by its covariance matrix of all three-second windows of imaged activity without stimuli. The response for each individual stimulus averaged across the three seconds was then also defined by a vector in neural space. This vector of each stimulus response was multiplied by the spontaneous covariance matrix to estimate the amount of overlap between each response and the spontaneous activity. For this analysis of the population, we used all cells that were active with dF/F0 > 1 and tuned with an R^2^>0.7. Overlap was calculated with the following equation ^45,81^:

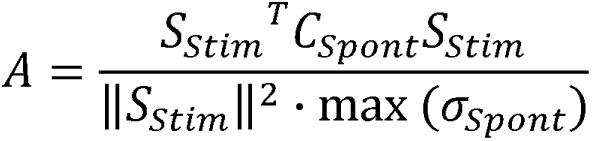

The neural response vector to each stimulus, *S*_Stirn_, was projected into the neural space that was observed during the spontaneous neural activity, defined by its covariance matrix, *c*_Spont_. We normalized by dividing by the largest singular value of the covariance matrix, *c*_Spont_. Each stimulus response was also normalized to a unit vector by dividing by the squared Euclidean norm of the *S*_Stirn_ to compare relative responses across cells rather than global activity levels. Thus, the amount of overlap could range from 0 to 1, with a maximum overlap of 1 when with *S*_Stirn_ perfectly aligns to the spontaneous neural dimension with the largest variance and 0 if *S*_Stirn_ aligns with a neural dimension with no spontaneous activity. For comparison among the four different grating pattern stimuli, we calculated the percentage of spontaneous activity that overlapped with each individual grating out of the total overlap across the four different patterns within a cycle. A two-way ANOVA with S1 vs. S2 and animal as the two factors for the 8 stimuli cycles was used to test for significant differences in the overlap between the pre-and post-altered experience imaging sessions for the four grating stimuli and to generate 95% confidence intervals.

### Statistical analysis

Statistical tests were performed using Prism 9, SPSS, or MATLAB. No statistical methods were used to predetermine sample sizes. The sample sizes are comparable to previous literature. Test for normal distribution was done with the Kolmogorov-Smirnov test. *p* < 0.05 was considered statistically significant. Sample sizes are reported in figure legends. Linear mixed effects model was used to compare nested data. Stimuli and mice were used as fixed and random factors, respectively. Samples were individual mice or neurons or stimulus cycles (indicated in legends). For obtaining 95% confidence intervals for similarity ratios, 1000 similarity values between two stimuli were obtained by bootstrapping with a replacement for the S1 and S2 data set, and the S2-S1 ratio was calculated. The bootstrapped ratios were sorted in ascending order, and 50th and 950th values were used as the lower and upper bound of the 95% confidence interval. Similarly, 100th and 900th values were taken as the lower and upper bound of a 90% confidence interval. Statistical procedures are two-sided and are listed in figure legends.

### Availability of materials and data

- The datasets generated during and/or analyzed during the current study are available from the the corresponding author on request.

## Acknowledgment

This work was supported by a grant from the National Institutes of Health (Grant No. RO1AG064067) to J.S. A.G.R was supported by research grant NIH-R00NS101127. We thank the members of the Subramanian lab for their comments on the manuscript.

## Author contributions

S.N. and J.S. designed the experiments. W.L.H, A.G.R, and J.S. performed data analysis. A.G.R. and J.S designed data analysis. J.S. wrote the paper and supervised the research.

